# Ultrashort optical-pin excitation for scattering brain imaging

**DOI:** 10.64898/2026.01.04.697592

**Authors:** Xiaobin Weng, Qiannan Song, Cihang Kong, Xin Dong, Qingliang Zhao, Jun Dong, Hongsen He

## Abstract

Imaging neural structures deep in brain tissue is central to understanding brain function, yet remains fundamentally limited by strong optical scattering and the requirement for accurate three-dimensional (3D) optical sectioning. Laser-scanning microscopy is a promising technique for brain imaging; however, maintaining excitation focus integrity in scattering media while preserving axial confinement poses a persistent photonic challenge. Here we introduce the optical pin, an ultrashort excitation regime engineered at the angular-spectrum level to address this limitation. By broadening the transverse angular bandwidth of a Bessel-type field while preserving its conical momentum-space architecture, the optical pin introduces a controlled longitudinal wave-vector spread that compresses the axial interference length to the micrometer scale, restoring Gaussian-like sectioning without sacrificing multi-angle interference. This excitation design yields substantially enhanced imaging performance, including ∼1.5-fold contrast improvement and ∼2.6-fold increased robustness to scattering. We validate the approach across transparent, scattering, and biological specimens, including bead phantoms, *C. elegans*, and mouse brain tissue. As a system-level excitation strategy, the optical pin is readily compatible with existing laser-scanning microscopy platforms and is particularly suited for scattering-limited brain imaging.

## Introduction

Laser-scanning multiphoton microscopy (MPM) has become a cornerstone for *in vivo* recordings of neuronal activity [1], capitalizing on nonlinear light–matter interactions to deliver intrinsic optical sectioning and reliable three-dimensional (3D) imaging [2-4]. The prevailing implementation relies on a Gaussian excitation beam, whose tight focus yields robust axial confinement under weakly scattering conditions [5-8]. However, as applications push toward deeper brain regions and more heterogeneous tissue, Gaussian-beam MPM increasingly encounters fundamental limitations: multiple scattering and specimen-induced aberrations degrade the focus, reduce contrast, and compromise both image quality and penetration depth [9, 10]. To overcome these challenges, numerous strategies have been explored, including high nonlinear excitation (e.g., three-photon) to ensure sufficient signal-to-noise ratio (SNR) in deep regions [11-13], adaptive optics to actively correct wavefront distortion [14, 15], or tissue clearing to reduce the scattering in the sample [16, 17]. While powerful, most approaches usually demand complex hardware, iterative calibration, increased computational load, or are confined to specialized samples. By contrast, engineering the focal field itself offers a straightforward, hardware-friendly route that integrates seamlessly with laser-scanning microscopes [18-20]. Representative examples include axially structured illumination [21, 22], continuous multifocal beam [23], droplet beam [20, 24], and various nondiffracting beams [25].

Among these options, the nondiffracting beams have attracted particular interest, especially the Bessel beam for volumetric laser-scanning microscopes [26-30]. Its sharply confined central lobe promotes high contrast in imaging; its nondiffracting behavior extends depth of field (EDOF); and its self-healing property renders the focus more resilient to perturbations such as scattering and partial occlusions. Yet the same axial elongation that underpins EDOF also collapses volumetric information into a 2D projection, erasing axial detail and restricting application primarily to sparse samples. This well-known trade-off motivates a natural question: can one retain the contrast and scattering resilience of Bessel beam while restoring Gaussian-like optical sectioning?

Here, we answer this question by introducing an ultrashort Bessel beam for laser-scanning microscopes that combine the strengths of both excitation paradigms. The key idea is to deliberately shorten the Bessel focus so that axial resolution is recovered, while preserving a compact central lobe and the multi-angle interference that underlies self-healing. This configuration (i) compresses the Bessel length to reinstate optical sectioning, (ii) further shrinks the central spot to enhance image contrast, and (iii) maintains the constructive interference of plane-wave components arriving from multiple angles, so that each axial position is formed by multiple routes, making the excitation less susceptible to scattering. Although the past two decades have seen extensive use of long Bessel beams across laser-scanning platforms, including coherent Raman scattering [31, 32], photoacoustic [33, 34], and multiphoton [28] microscopy, investigations of short-focus Bessel beams remain scarce.

In this work, we propose the concept of ultrashort nondiffracting beam for simultaneous optical-sectioning and robust-scattering imaging, with the added benefit of high contrast. Opposite to the conventional axially elongated beam (known as an “optical needle” [35, 36]), this ultrashort Bessel beam is termed an “optical pin”. We detail a practical procedure to produce the optical pin and validate its performance in a laser-scanning MPM platform. Experimentally, we shorten the Bessel length from 48 to 5 μm, achieving Gaussian-like axial resolution while delivering ∼44% higher image contrast and ∼4.1 dB greater scattering tolerance (compared to a Gaussian focus). We compare imaging performance between transparent and scattering bead phantoms, and further demonstrate its applicability through biological imaging of *C. elegans* and mouse brain neurons. These results establish the optical pin as a promising, reliable excitation mode for high-contrast, scattering-resilient 3D MPM, and can be readily applied to other laser-scanning microscopy platforms.

## Results

### Design of optical pin

We first outline the design rationale of the optical pin in the context of limitations faced by conventional Gaussian and long Bessel beams [Fig. 1(a)]. Bessel beams are typically utilized in laser-scanning microscopy for their EDOF, enabled by their nondiffracting propagation, to facilitate rapid volumetric scans. However, if the nondiffracting length of a Bessel beam can be deliberately shortened to the scale of a Gaussian focal depth, it would attain optical sectioning capability comparable to a Gaussian beam, thereby enabling reliable true 3D imaging. Importantly, in this shortening process of the Bessel beam, its inherent self-healing and multi-angle interference properties are preserved, providing additional advantages in scattering tolerance and high-contrast imaging over a Gaussian beam. In analogy to the “optical needle” referring to a long Bessel beam, we use the term “optical pin” to describe the ultrashort Bessel beam configuration [Fig. 1(b)].

**Fig. 1.**
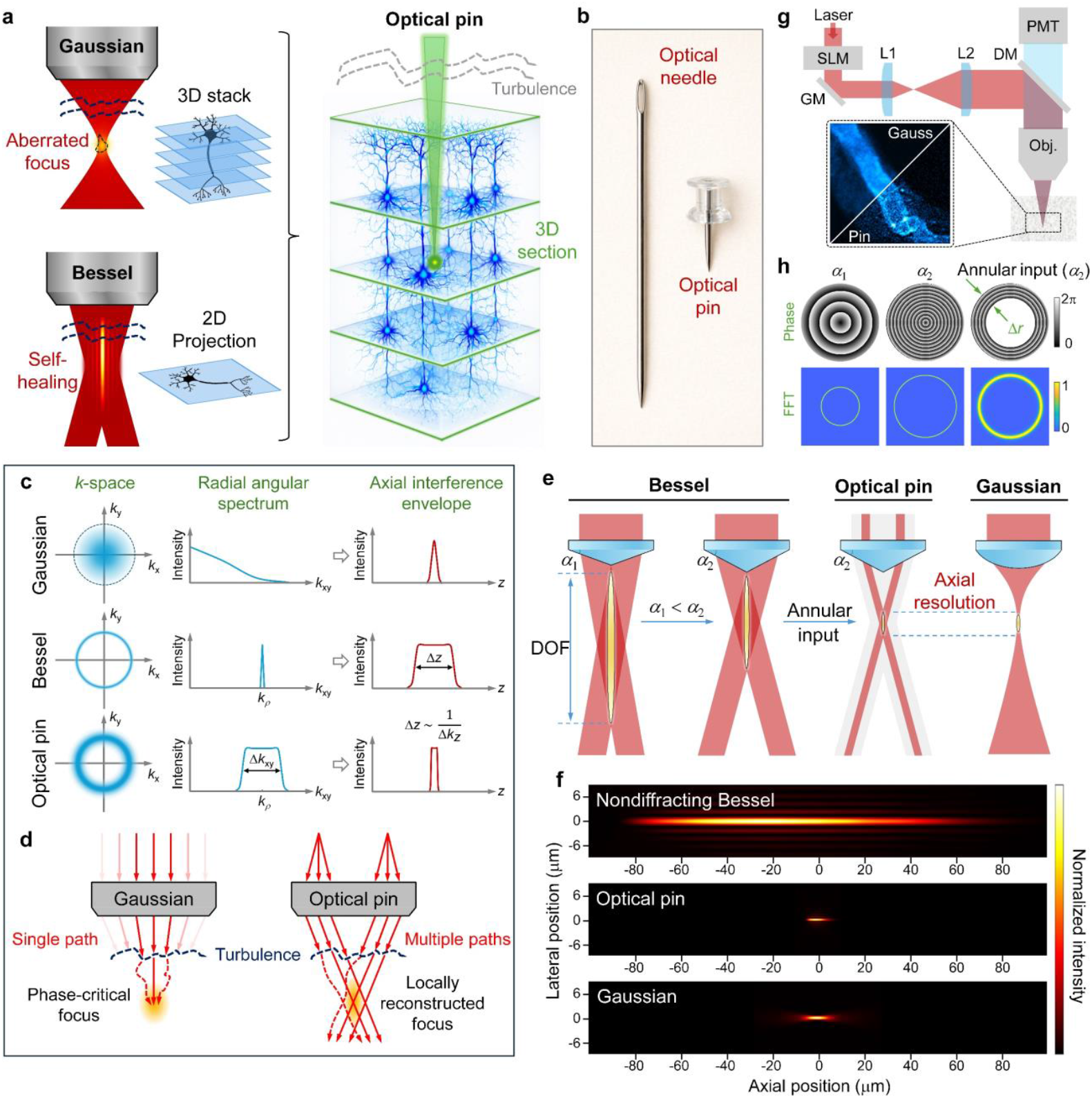
Design and utilization of optical pin. (a) Comparison of conventional Gaussian and Bessel beams in laser-scanning microscopes. The Gaussian beam holds the capacity for optical sectioning, but has low tolerance for tissue turbulence; while the conventional Bessel beam has self-healing feature in tissues, but projects volumetric information into a 2D image. The optical pin combines the merits of both beams for optical sectioning, anti-scattering, and high contrast. (b) Concept of the “optical needle” and “optical pin” for microscopy. (c) Frequency-space interpretation of Gaussian, Bessel, and optical-pin excitation. Column 1 shows their 2D angular spectra, with a central distribution for the Gaussian and thin/broadened annuli for the Bessel/optical pin. Column 2 presents the corresponding radial spectra, highlighting low-frequency Gaussian excitation and narrow/broadened annular distributions for the Bessel/optical pin. Column 3 illustrates the resulting axial profiles: a finite Gaussian focus, an extended nondiffracting Bessel region, and a compressed axial confinement for the optical pin. (d) Schematic comparison of scattering robustness. Gaussian excitation relies on a single phase-critical path and forms a distorted focus under turbulence, whereas the optical pin combines multiple angular paths to produce phase-tolerant, locally reconstructed foci that preserve focal integrity under scattering. (e) Approaches to creating the optical pin. (f) Numerical simulations of the three beams. (g) Experimental configuration of laser-scanning microscopes under optical-pin excitation. The zoomed image illustrates the contrast enhancement with the optical pin in MPM. (h) Phase patterns on the SLM and the corresponding FFT images for Bessel beams and optical pin. **(Visualization 1)**

Fig. 1(c) provides a frequency-domain interpretation of axial compression through angular-spectrum engineering. As shown in Column 1, Gaussian, conventional Bessel, and optical-pin excitation exhibit distinct transverse wave-vector distributions in the back focal plane: the Gaussian beam concentrates near the optical axis, the Bessel beam forms a thin annulus at a fixed cone angle, and the optical pin broadens this annular support while preserving a non-zero transverse momentum. Radial representations of the spectra (Column 2) clarify the role of angular bandwidth. Although the Gaussian beam spans a finite transverse bandwidth, its spectral weight is confined to low spatial frequencies, whereas the Bessel beam enforces a narrow distribution around a specific *k*_*ρ*_. By deliberately expanding the annular bandwidth Δ*k*_*xy*_, the optical pin introduces a finite longitudinal wave-vector spread Δ*k*_*z*_, which compresses the axial interference envelope, as illustrated in Column 3. Importantly, the finite axial extent of a Gaussian focus does not originate from a large Δ*k*_*z*_, since most components satisfy *k*_*z*_ ≈ *k*, but from the absence of a structural constraint that locks longitudinal phase evolution, leading to rapid dephasing away from focus and geometry-limited axial confinement. In contrast, a conventional Bessel beam enforces a fixed cone angle that locks across angular components, resulting in an extended nondiffracting region. The optical pin operates between these regimes: it retains the conical momentum structure while introducing a controlled angular bandwidth, yielding a non-zero Δ*k*_*z*_ and a shortened axial interference length. Quantitative relationships among Δ*k*_*xy*_, Δ*k*_*z*_, and the resulting axial extent Δ*z* are given in the Methods. Fig. 1(d) illustrates the contrasting responses of Gaussian and optical-pin excitation to scattering-induced phase perturbations. Gaussian focusing relies on a single near-axis propagation channel and globally phase-matched interference, making the focal peak highly sensitive to turbulence and scattering. In contrast, the optical pin forms its focus through multiple angular paths distributed over a conical momentum support. Although individual components may be perturbed, a subset remains locally phase-matched, enabling statistically robust focal reconstruction. This angular redundancy renders the optical-pin focus intrinsically phase-tolerant, such that scattering robustness emerges from momentum-space architecture rather than from an extended nondiffracting length.

The frequency-domain analysis in Figs. 1(c, d) indicates that axial compression and scattering robustness arise from engineering the transverse angular spectrum, specifically by broadening the annular support at a finite cone angle. In practice, this spectral condition is not achieved by a single experimental parameter, but by the combined control of the annular illumination geometry and the cone angle of the axicon or objective. These two degrees of freedom map directly onto the width and the central position of the transverse *k*-space distribution, respectively, and together determine the longitudinal wave-vector spread and the resulting axial interference length. Fig. 1(e) illustrates the approaches to shorten the axial extent of a Bessel excitation. The nondiffracting range is described as

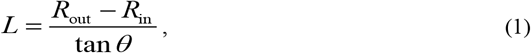

where *R*_out_ and *R*_in_ are the effective outer and inner radius of the incident beam on the conical surface of the axicon, respectively [Fig. S2]. For a conventional Gaussian input, *R*_in_ = 0. The axicon generates a conical wave with a deflection angle *θ* that is given by

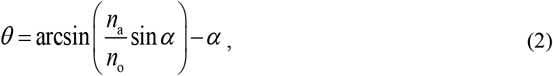

where *α* is the physical angle of the axicon. *n*_a_ and *n*_o_ are the refractive indices of the axicon and the surrounding medium, respectively. In MPM (*m*-photon excitation), the effective length also depends on the axial excitation nonlinearity, thus the Bessel length in MPM is

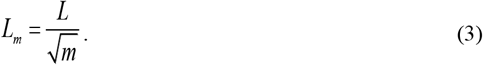

According to Eqs. (1-3), the Bessel length is mainly determined by the thickness Δ*r* of the incident annular beam and the physical angle *α* of the axicon: a thinner ring beam shines into an axicon with a larger physical angle, producing a shorter nondiffracting region. We denote this shortened nondiffracting length in MPM as *L*_m_, which corresponds to the axial resolution achieved by the ultrashort Bessel beam under *m*-photon excitation. As illustrated in Fig. 1(e), the Bessel length is decreased when the physical angle of the axicon increases from *α*_1_ to *α*_2_, and it can be further shortened to the scale of a Gaussian focal depth by utilizing the annular beam input, achieving the same axial resolution.

For the lateral resolution of the optical pin in MPM, its multiphoton signal can be expressed as

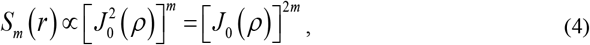

where *J*_0_(*ρ*) is the zero-order Bessel function of the first kind, determining the position of the main lobe and concentric rings. *ρ* = *k*_o_*r*sin*θ* and *k*_o_ = 2π*n*_o_/*λ*, where *λ* is the laser wavelength in the air and *r* is the lateral coordinate. The lateral resolution is defined as the full width at half maximum (FWHM) of the Bessel main lobe under multiphoton excitation. Thus, letting [*J*_0_(*ρ*_1/2_)]^2*m*^ = 1/2, the resolution can be expressed in paraxial approximation:

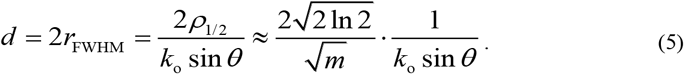

According to Eq. (5), the lateral resolution of the Bessel beam is improved by a large physical angle and high-order photon excitation. Based on the above equations, we can find that shortening the Bessel length will improve the overall 3D resolution, and the use of multiphoton excitation further enhances this effect. This feature endows the optical pin with a high-contrast optical sectioning capability in MPM. Numerical simulations confirm these design considerations [Fig. 1(f)]. The optical pin exhibits significantly improved 3D resolution compared to a typical long (nondiffracting) Bessel beam. At the same time, it maintains a thinner lateral main lobe and still retains a nondiffracting propagation character at defocused planes (both axial ends), in contrast to a Gaussian beam that diverges and suffers intensity loss away from focus [Fig. S3].

Fig. 1(g) illustrates the implementation of the optical pin in a standard laser-scanning MPM platform. A phase-only spatial light modulator (SLM) is placed at the conjugate pupil plane, enabling programmable switching between a lens phase for Gaussian focusing and an axicon phase for Bessel excitation within the same optical path. As shown in Fig. 1(h), the effective axicon physical angle (*α*_1_ and *α*_2_) is digitally controlled by the conical phase, thereby tuning the transverse *k*-space distribution and the corresponding Bessel length. The optical pin is realized by further imposing an annular condition on the axicon phase, removing the central region so that only a thin ring contributes to the conical interference. This configuration implements the two-step shortening strategy in Fig. 1(e), compressing the axial extent to the micrometer scale while preserving the multi-angle interference responsible for self-healing. The corresponding fast Fourier transforms (FFTs) represent the intensity distributions at the back focal plane of the objective and thus the encoded transverse *k*-space spectra. Increasing the axicon angle from *α*_1_ to *α*_2_ enlarges the radius of the annular pattern while keeping a similar ring thickness, indicating higher transverse wave vectors with comparable angular bandwidth. Removing the central region of the *α*_2_ phase produces a thicker annulus at the same radius, corresponding to a broader angular spectrum that shortens the axial interference length and generates the optical pin.

### Optical pin generation in MPM

Following the above design, we experimentally generated a series of Bessel beams and characterized them in a MPM setup. The axial images and intensity profiles are shown in Figs. 2(a-c). We gradually shortened the beam length from 48 to 5 μm by adjusting the axicon angle and shape of the incident beam. Using a full (solid) Gaussian illumination on the SLM, the Bessel length could be tuned from 48, 37, to 29 μm by directly increasing the axicon angle, corresponding to effective objective lens NAs of approximately 0.55, 0.8, and 0.9, respectively. Beyond ∼29 μm, the length cannot be further shortened, as the Bessel beam has fully filled the back aperture of the objective (0.9 NA). Then, the incident beam is switched from the solid beam to the annular beam, achieved by deflecting the central part of the beam on the SLM [see phase patterns in Fig. S4]. By decreasing the thickness of the effective reflection region, the Bessel length is further shortened from 13 to 5 μm. For comparison, we also generated a conventional Gaussian focal spot with an axial length of ∼5 μm, matching the 5-μm optical pin [Figs. 2(a, c)].

**Fig. 2.**
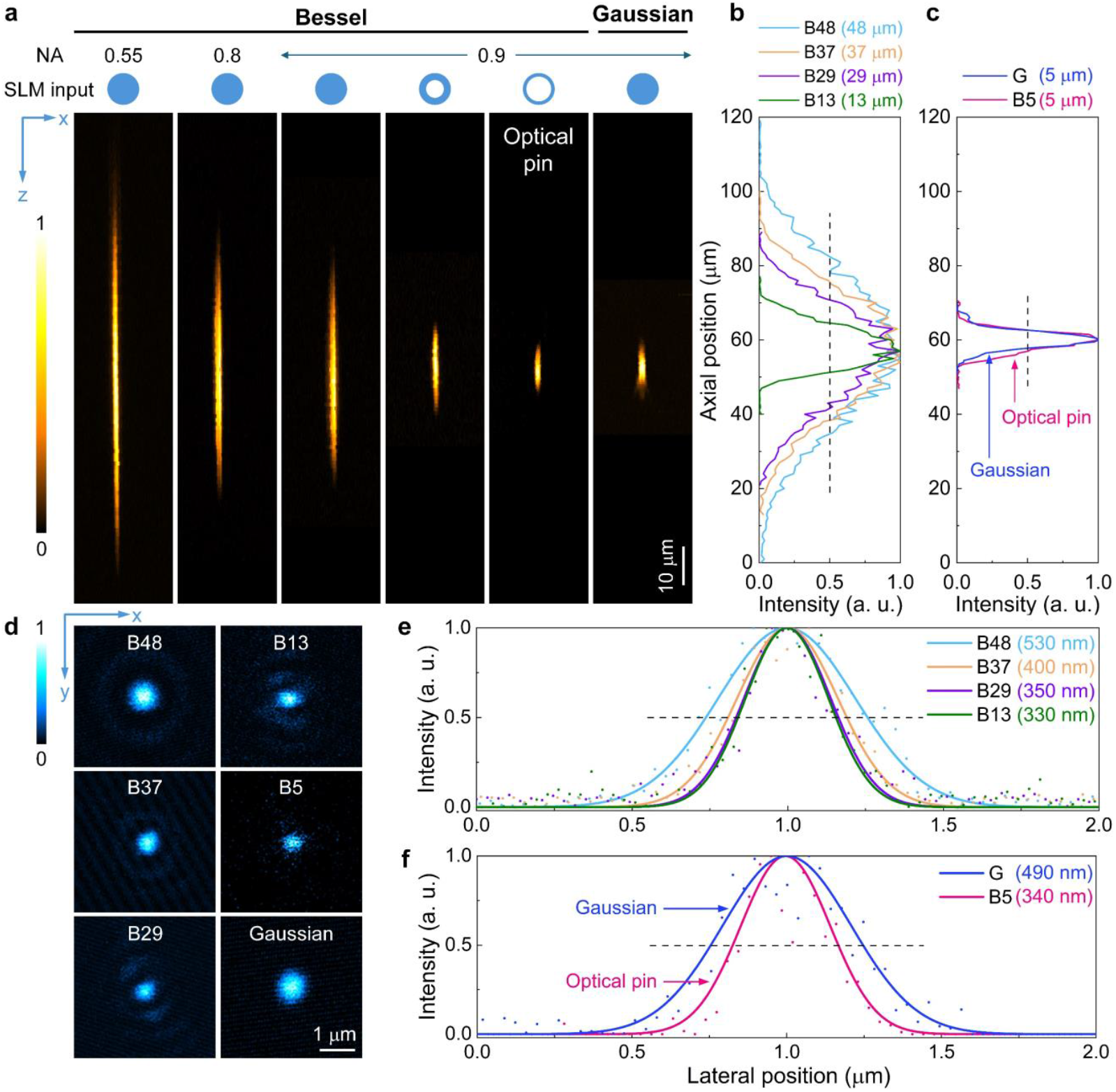
Generation of optical pin in MPM. (a) PSFs of the Bessel beams in the *x*-*z* plane with different lengths, ranging from 48, 37, 29, 13, to 5 μm (optical pin), and the Gaussian PSF (5 μm). NA: the effective NA of the objective lens, depending on the beam size at the back focal plane of the objective. SLM input: the shape of the incident beam on the SLM. (b, c) Axial intensity profiles of the Bessel, optical pin, and Gaussian beams, corresponding to (a). (d) PSFs of the Bessel, optical pin, and Gaussian beams in the *x*-*y* plane. (e, f) Lateral intensity profiles of the three beams, corresponding to (d).

The lateral PSFs and intensity profiles for these beams are shown in Figs. 2(d–f). As the effective NA increased from 0.55 to 0.9 (solid-beam cases), the Bessel central spot diameter (FWHM) decreased from ∼530 nm to ∼350 nm, owing to the inclusion of higher spatial frequency components under larger axicon angles. With the annular beam input at 0.9 NA, the main spot size remained approximately 330 - 350 nm even as the axial length was reduced from 13 to 5 μm. In comparison, the Gaussian PSF at the same 0.9 NA had a FWHM of ∼490 nm. Thus, the optical pin central lobe was ∼30.6% narrower than the Gaussian, indicating a significant improvement in the lateral image contrast. Because the spatial-frequency support set by the objective NA is unchanged, the smaller central-lobe FWHM reflects contrast enhancement via energy redistribution rather than a genuine increase in lateral resolution. These measurements demonstrate that by compressing the Bessel length to ∼5 μm, one can achieve Gaussian-like axial confinement while maintaining a smaller lateral spot than a Gaussian beam at the same NA. Notably, this axial shortening was accomplished with minimal change in the lateral spot size, highlighting that the optical pin decouples axial and lateral resolution to a large extent. This ability to independently tune axial confinement is unique to Bessel-type beams.

### Optical pin imaging in scattering samples

To evaluate the high-contrast and anti-scattering features under the optical-pin excitation, we applied it to image fluorescent microspheres in both transparent and scattering media [Fig. 3]. The transparent sample consisted of 1-μm fluorescent beads in pure water, whereas the scattering sample contained the same beads mixed with 1-μm TiO_2_ nanoparticles to introduce controlled light scattering [see Fig. S5 for details]. As shown in Fig. 3(a), under transparent conditions, the optical pin produces bead images with sharper edges and higher contrast than the Gaussian beam. This is confirmed by line profile measurements in representative regions, which show contrast enhancements of ∼2.1 and ∼2.9 dB for the optical pin relative to the Gaussian focus [Fig. 3(b)]. In the scattering sample [Fig. 3(c)], the optical pin demonstrates even greater improvements in image quality. The optical-pin-excited image exhibits brighter bead signals and a lower background haze compared to the Gaussian-excited image, indicating a higher SNR. This superior performance is attributed to the Bessel self-healing nature, which is retained by the optical pin regardless of its shortened length. The spatial frequency spectra [Fig. 3(d)] of the captured images reflect this difference: the optical pin preserves higher spatial frequency content than the Gaussian in the scattering case, consistent with improved contrast. Detailed comparisons of three example regions [Fig. 3(e)] further illustrate that beads are more clearly resolved with the optical pin excitation. A quantitative SNR analysis of the scattering experiment [Fig. 3(f)] indicates a ∼4.1 dB improvement in SNR for the optical pin over the Gaussian. These results demonstrate that the advantages of optical pin are even more pronounced in a scattering environment, confirming its strong robustness to tissue-like turbidity. This behavior is consistent with previous observations that Bessel beams tend to maintain focus integrity better than Gaussian beams in heterogeneous media [37].

**Fig. 3.**
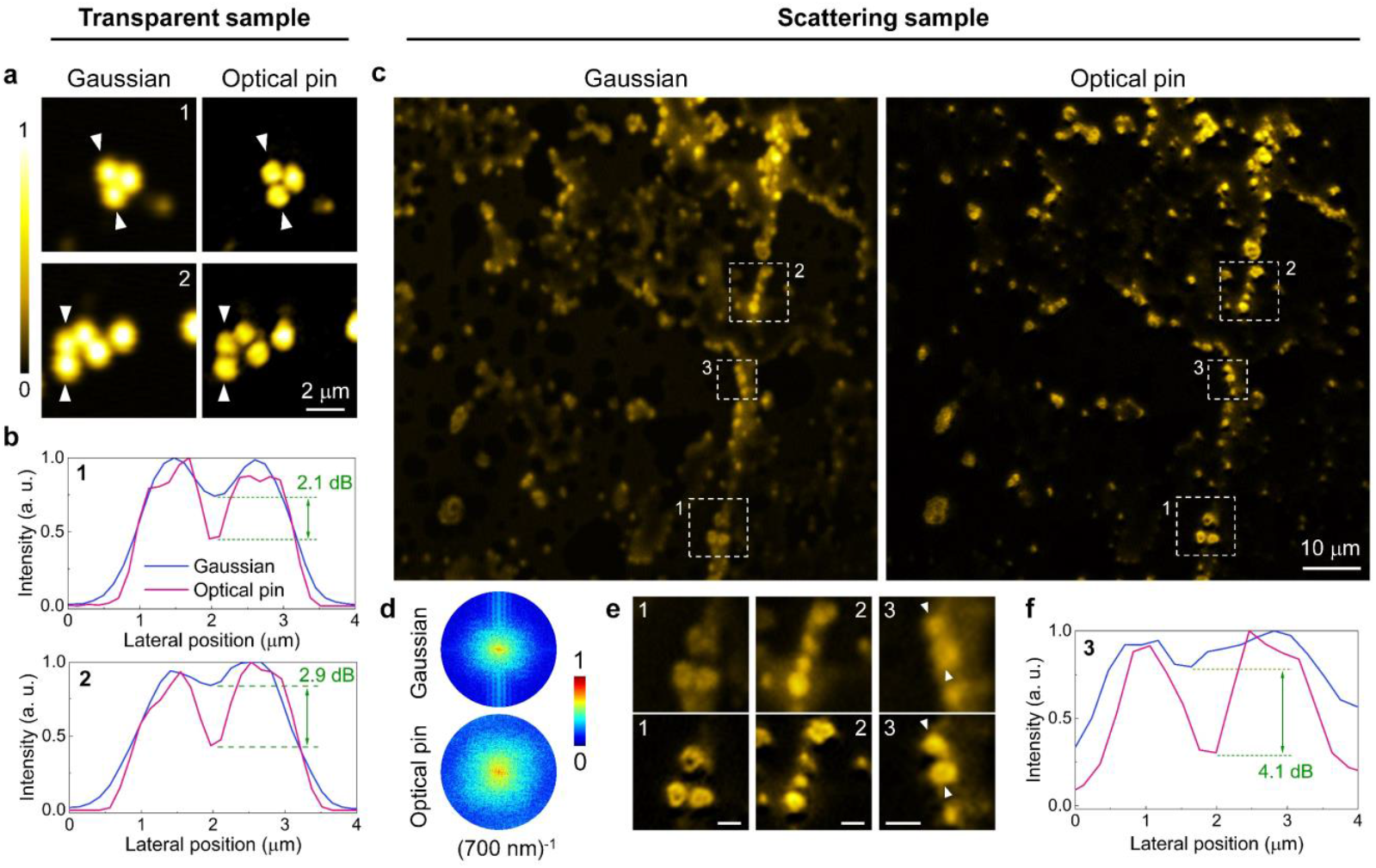
Optical-pin MPM for transparent and scattering samples. (a) Comparison of Gaussian and optical-pin images for fluorescent beads in transparent medium (water). (b) Line profile comparisons for typical regions, indicated by the white arrows in (a). (c) Comparison of Gaussian and optical-pin images for fluorescent beads in scattering medium (mixed with 1-μm TiO_2_ particles). The comparison of the sample photos is shown in Fig. S5 of the Supplementary. (d) Spatial frequency domain of the images in (c). The edge of the frequency circles is located at (700 nm)^-1^. (e) Zoomed regions in (c). Scale bar: 2 μm. (f) Line profile comparison for region 3, indicated by the while arrows in (e).

### High-contrast imaging for *C*. elegans

To showcase the optical pin performance in a biological tissue context, we next imaged fluorescently labeled *C. elegans* samples with both Gaussian and optical-pin excitation [Fig. 4]. In these specimens, the intestinal lumen was labeled via a fluorescently tagged actin (ACT-5) expressed in the microvilli of the intestinal cells. As shown in Figs. 4(a, d), the optical pin image maintains all the structural details visible in the Gaussian-beam image, but with noticeably sharper delineation of the intestinal wall boundaries. This improvement is confirmed by the zoomed-in regions [Figs. 4(b, e)] and their intensity line profiles [Figs. 4(c, f)]. The optical pin yields steeper intensity transitions at the lumen edges, indicating enhanced contrast and edge definition. Importantly, this higher contrast is achieved without an increase in imaging power or acquisition time relative to the Gaussian case (see Methods). The ability to better resolve fine anatomical details under identical conditions underscores the practical advantage of the optical pin for biological imaging: it can reveal subtle structural features more clearly, which could facilitate more accurate morphological or functional observations in living specimens.

**Fig. 4.**
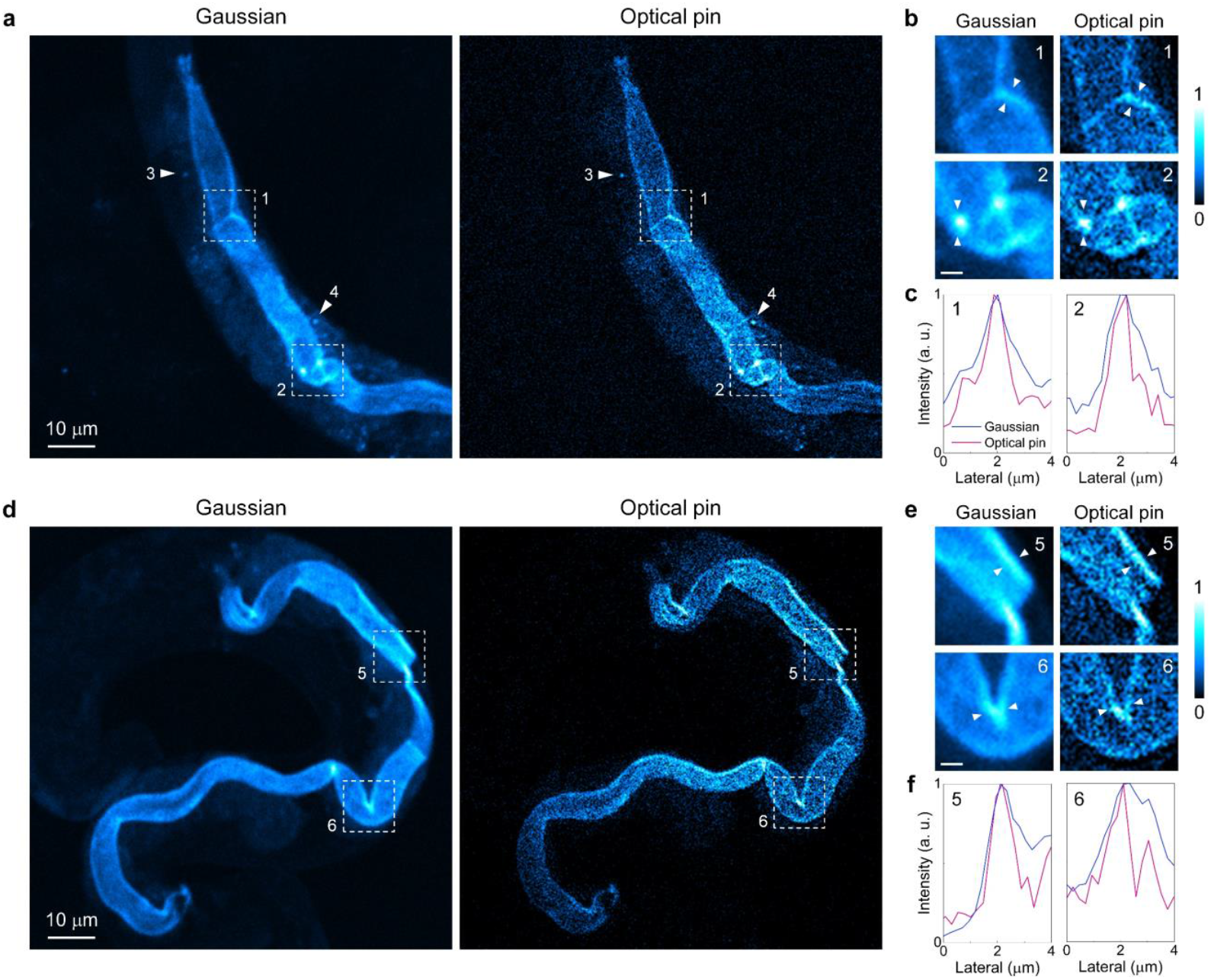
High-contrast imaging of optical-pin MPM for *C*. elegans. (a, d) Gaussian and optical-pin images for *C*. elegans. (b, e) Zoomed regions in (a, d) for comparison. Scale bar: 2 μm. (c, f) Line profiles of typical positions in (b, e).

### Optical sectioning and robust scattering in mouse brain tissues

Having established the anti-scattering and high-contrast benefits of the optical pin in controlled samples and single-layer *C. elegans*, we further validated its performance in a densely labeled, strongly scattering biological specimen. We imaged a thick mouse brain section (100 μm) containing neurons expressing tdTomato, using both the optical pin and a standard Gaussian beam for comparison. A 3D stack of MPM images was acquired by axially scanning the focus through the tissue, and the resulting volume was projected into a single 2D image with depth encoded by color [Fig. 5(a)]. Strikingly, the optical pin excitation reveals more neuronal structures in this thick tissue than the Gaussian excitation. In the optical-pin image, neuron processes appear sharper and additional fine branches are visible in several regions that are barely discernible or missing in the Gaussian image, where examples are indicated by dashed boxes in Fig. 5(a). The FFT in Fig. 5(d) also illustrates more high-frequency components in the optical-pin image. This dramatic improvement can be attributed to two factors. First, for the optical pin, its smaller main lobe (tighter focus) provides inherently higher contrast, making faint structures stand out more clearly. Second, its multi-route interference focus is less sensitive to sample-induced aberrations and scattering, resulting in fewer shadow artifacts than the single-path, plane-wave Gaussian focus. As a result, the optical pin can penetrate deeper with maintained resolution, whereas the Gaussian focus degrades more quickly with depth.

**Fig. 5.**
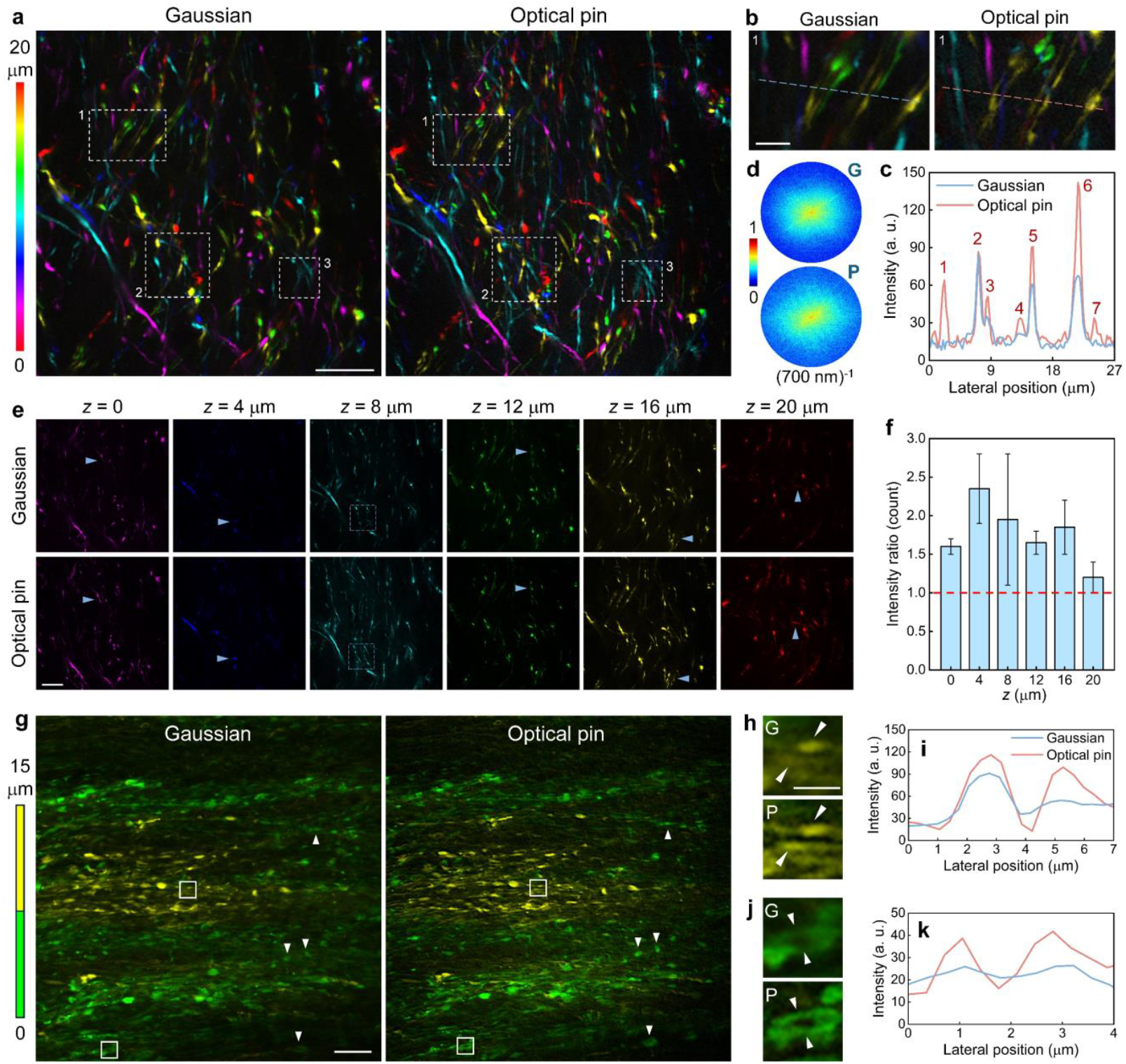
Simultaneous optical-sectioning and robust-scattering brain imaging with high contrast via optical pin. (a) Comparison of the 3D neuron images obtained by the Gaussian beam and optical pin. The imaging depth is color-coded. (b) Zoomed regions in (a). (c) Intensity profile comparison of the neurons for the two beams. (d) FFT comparison for (a). The edge of the frequency circles is located at (700 nm)^-1^. (e) Plane-by-plane comparison between the Gaussian beam and optical pin. Depth located at 0, 4, 8, 12, 16, and 20 μm of brain tissue. The arrows point to the positions for comparison. (f) The fluorescence intensity ratio of the same neuron structure from the optical pin to the Gaussian beam at varying depths. (g) Comparison of the neurons in dense regions of the brain under the excitations of the two beams. (h-k) Zoomed positions (h, j) and line profiles (i, k) from (g) for comparison. Scale bar: 20 μm for (a, e, and g) and 5 μm for (b, h, and j). **(Visualization 2)**

Fig. 5(b) highlights a magnified region from Fig. 5(a), comparing the two excitation modes in detail. Thin processes of the neurons that are almost lost in the Gaussian image are much better resolved with the optical pin. The corresponding intensity profiles [Fig. 5(c)] show that what appears as a single broad peak (or remains unresolved) in the Gaussian case is resolved into multiple distinct peaks with the optical pin, including several low-intensity neurites that only the optical pin could reliably excite. A plane-by-plane comparison through the image stack [Fig. 5(e)] further illustrates the difference: at each depth, the optical pin produces a higher SNR of the neurons than the Gaussian, as indicated by the white arrows highlighting example neuron positions. To quantify this improvement across depth, we calculated the fluorescence intensity ratio for corresponding neuronal structures imaged with the optical pin versus the Gaussian beam [Fig. 5(f)]. The fluorescence signal (for the same structure at each depth) was normalized to the maximum signal in its respective imaging depth. We found that the optical pin provided higher fluorescence intensity for the same structures at all depths examined (enhancement ranging from ∼1.2-fold to ∼2.4-fold), confirming the robust scattering performance of the optical pin throughout the 3D volume of the tissue. Finally, we assessed the optical-pin imaging ability in a densely labeled brain region to ensure that its benefits hold even amid overlapping structures. Two example imaging planes (at depths 0 and 15 μm) in a fiber-rich brain region are shown in Fig. 5(g) for both excitation types. Both the Gaussian and optical pin can excite the same overall structures and allow depth discrimination, but the optical pin reveals more fine details within this dense network, including small neural processes that are much less apparent in the Gaussian images. Zoomed-in subregions [Figs. 5(h, j)] and their line profile plots [Figs. 5(i, k)] confirm that the optical pin uncovers structures that produce either relatively weak or no signal under Gaussian excitation. Even in this complex, high-density environment, the optical pin provides clearer separation of adjacent structures and higher image contrast.

## Discussion

In laser-scanning microscopy, the characteristics of the focused beam are critical to imaging quality, including contrast, penetration depth, and optical sectioning capability. The conventional tightly focused Gaussian beam provides strong axial confinement but suffers from degradation due to scattering-induced aberrations as it propagates in tissue [37]. The long nondiffracting Bessel beam, on the other hand, can resist scattering through its self-healing propagation, but it lacks axial resolution and true sectioning ability [27]. In this work, we combine the merits of these two beam types into a new excitation mode, termed the optical pin (as opposed to the extended optical-needle beam [36]). The optical pin is designed to simultaneously retain the high contrast and scattering resilience of a Bessel beam while restoring Gaussian-like optical sectioning.

The optical pin is created by extremely shortening the length of a conventional Bessel beam. Two ways are applied together to compress the axial nondiffracting range: using large angel axicon and an annular beam input [Fig. 2(a)]. The former tightly interferes the conical wavefronts to push the Bessel main lobe further small for high imaging contrast, while the latter directly cancels the Bessel length for achieving 3D sectional capacity. Based on the two steps, the US Bessel beam is created, which is also a significant complement to the Bessel family. Although the US Bessel beam is short, it still maintains the self-healing advantage. The self-healing property of an ultrashort Bessel beam originates from its conical momentum-space structure rather than its axial non-diffracting length [30]. Although axial interference is truncated, the transverse wave-vector redundancy is preserved, allowing the central lobe to be reconstructed through angularly distributed off-axis components. In contrast, Gaussian beams localize phase coherence at a single focal plane and lack angularly distributed transverse momentum components; once the focal region is perturbed, the beam cannot reconstruct its central lobe during subsequent propagation [10]. The scattering-robust nature of the optical pin is evidenced by our imaging of the scattering bead phantom, where the optical pin maintained much higher contrast than the Gaussian focus under identical conditions [Fig. 3(c)].

The high imaging contrast of the optical pin comes from the smaller Bessel central lobe than the Gaussian spot. Conventional long Bessel beams, as commonly used in volumetric laser-scanning microscopes, already exhibit a substantially narrower main lobe than Gaussian beams [28], despite their extended axial extent. This advantage originates from the conical angular spectrum of Bessel beams, which concentrates transverse momentum at a well-defined wave-vector magnitude and sustains a tightly confined central lobe through continuous angular interference rather than single-plane focusing. In our work, this intrinsic lateral confinement is further enhanced by deliberately using a large-physical-angle excitation. As a result, the transverse wave-vector content contributing to the central lobe is further sharpened, leading to an additional reduction of the main-lobe width [Fig. 2(d)]. The high-contrast feature is confirmed by the single-layer *C. elegans* sample [Fig. 4] and the thick brain tissue [Fig. 5]. Our approach therefore does not merely preserve the lateral advantage of long Bessel beams, but actively amplifies it, achieving a smaller and more robust excitation profile than both Gaussian focusing and previously reported long Bessel configurations.

Beyond scattering robustness and high-contrast excitation, an important advantage of the ultrashort Bessel beam demonstrated here is that it endows conventional non–optically sectioned beams with intrinsic optical sectioning capability. In our experiments, the axial extent of the ultrashort Bessel beam is approximately 5 μm [Fig. 2(c)]; however, this length is not a fundamental limit and can be further reduced through straightforward optical scaling. First, employing objectives with a larger NA increases the accessible transverse wave-vector components, thereby shortening the axial interference length while simultaneously improving lateral confinement and contrast. For example, replacing the 0.9-NA objective used here with a high-NA objective (e.g., NA 1.3) is expected to reduce the Bessel length to the 1-3 μm range, and simultaneously a further smaller main spot at ∼200-300 nm theoretically. Second, a thinner annular illumination further broadens the angular spectrum, leading to an additional reduction of the axial extent without necessarily altering the lateral profile. Importantly, this behavior highlights a key distinction from Gaussian focusing, where lateral and axial dimensions are intrinsically coupled. In contrast, Bessel excitation allows axial confinement to be tuned largely independently of lateral resolution [Figs. 2(d-f)]. In the present implementation, the annulus thickness is constrained by the finite pixel size and damage threshold of the SLM, which limits the available excitation energy for very thin rings. This limitation can be readily overcome by alternative implementations, such as using an annular or thin-solid input of high energy to a single axicon [see Fig. S1 in Supplementary], offering a simple and scalable route toward even shorter and more intense optical-pin excitation.

Taken together, the optical pin demonstrated in this work simultaneously combines three key advantages within a single excitation scheme: high contrast, scattering robustness, and intrinsic optical sectioning. It is a capability that is not jointly available in existing beam configurations. Conventional approaches typically address these requirements separately and usually at the cost of increased system complexity. For instance, high-contrast imaging can be achieved through three-photon excitation [12], temporal focusing [38], or computational post-processing [6], based on additional hardware support or extensive training data. Scattering robustness, on the other hand, is most pursued using adaptive optics [14], usually relying on feedback loops, guide stars, or iterative optimization. Compared to depth-resolved strategies [39, 40], the optical-pin excitation provides 3D sectioning directly at the excitation level, without additional computation or hardware complexity. Moreover, many state-of-the-art volumetric imaging strategies are fundamentally built upon engineered Gaussian beams, including the reverberation loop [7], light beads [8], mirror-based axial scanning [41], and compressive multiplane imaging [21]. Replacing Gaussian excitation with an optical pin offers a fundamentally improved excitation field across such modalities. Owing to its strong resistance to scattering and precise axial confinement, this approach is particularly well suited for brain imaging, where highly scattering tissue and the need for accurate layer-specific interrogation coexist [2]. Finally, not restricted to MPM, this concept can be readily extended to other Gaussian-based laser-scanning modalities, such as photoacoustic [34] and coherent Raman microscopy [31]. This suggests a broad applicability of the optical pin across the photonics imaging field, providing a simple yet powerful strategy to improve imaging performance in complex, scattering samples.

## Methods and materials

### Relation between transverse angular bandwidth and axial confinement

The excitation field near focus is described using the angular spectrum formalism. In a homogeneous medium, each plane-wave component satisfies the dispersion relation

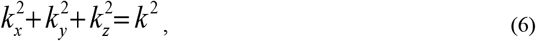

where *k* = 2π*n*/*λ* is the wave number in the medium. Defining the transverse wave vector magnitude

as 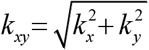, the longitudinal component is given by

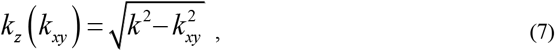

For a Bessel-type excitation, the transverse angular spectrum is concentrated around a finite cone angle, corresponding to a ring centered at *k*_*ρ*_ = *k*sin*θ*, where *θ* is the effective cone angle determined by the axicon or the NA of the objective. In experiments, increasing the axicon physical angle or using a higher-NA objective shifts the angular spectrum to larger *k*_*ρ*_, thereby increasing the sensitivity of *k*_*z*_ to variations in *k*_*xy*_.

Introducing annular illumination with a finite radial thickness broadens the transverse angular bandwidth Δ*k*_*xy*_ around *k*_*ρ*_· For a sufficiently narrow bandwidth (Δ*k*_*xy*_ << *k*_*ρ*_), the resulting spread of longitudinal wave vectors can be estimated by a first-order expansion,

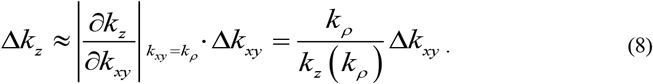

The axial interference length Δ*z* is then determined by the dephasing of longitudinal components and can be estimated by the coherence condition

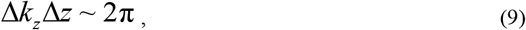

yielding Δ*z* ∼ 2π/ Δ*k*_*z*_.

This relation highlights the complementary roles of the two experimental controls employed here: annular illumination directly increases the transverse angular bandwidth Δ*k*_*xy*_, while a larger axicon angle or higher objective NA increases the central transverse wave vector *k*_*ρ*_. Together, these effects enhance the longitudinal wave-vector spread Δ*k*_*z*_ and compress the axial interference length, enabling the ultrashort Bessel regime termed the optical pin.

### Experimental setup

Fig. S1 shows the experimental setup for the optical-pin-excited laser-scanning MPM. The excitation light was from a mode-locked fiber laser (1030 nm, 19 MHz, and 180 fs) for two-photon excitation. The optical pin was generated by the SLM (HDSLM80R Plus, UPOLabs). The focal lengths of the lens are *f*_1_ = 200 mm, *f*_2_ = 200 mm, *f*_3_ = 100 mm, *f*_4_ = 100 mm, and *f*_5_ = 150 mm. A water-immersion objective (60×, 0.9 NA, LUMPLFL60XW, Olympus) was used to focus the excitation light into the sample. The *x*-*y* laser scanning was achieved by a pair of GMs (SG1150, Sino-Galvo Technology). The excited fluorescence signal was collected in the epi direction and detected by a photomultiplier tube (PMT, H11901-20, Hamamatsu). The residual excitation beam was cleaned up by two shortpass filters (OFE1SP-750, JCOPTIX) before the PMT. The electrical signal from the PMT was subsequently digitized by an analog-to-digital (A/D) converter card (PCI-6110, National Instruments, 5 MS/sec). The control of the entire imaging system, including laser scanning, data collection, and image reconstruction, was realized by a MATLAB-based multifunctional program.

Imaging parameters: For bead phantoms and *C. elegans* imaging [Figs. (3, 4)], Gaussian and optical-pin excitation were operated at the same effective post-objective power. Bead images were acquired at ∼13 mW, with a frame size of 512 × 512 pixels and a scanning speed of 0.43 Hz. *C. elegans* images were acquired at ∼27 mW, 512 × 512 pixels, and 0.2 Hz. For thick neuronal tissue imaging [Fig. 5], excitation power followed the commonly adopted practice in Bessel-based laser-scanning microscopy [19, 20, 24, 28]. Gaussian excitation was operated at ∼25 mW and optical-pin excitation used ∼95 mW, acquired with 512 × 512 pixels at a scanning speed of 0.15 Hz. No photobleaching or photodamage was observed under these conditions. The side lobes of the zeroth-order Bessel beams were cancelled by the corresponding third-order ones [26].

### Sample preparation

Fluorescence microspheres: Fluorescence beads (1 μm, F8819, Life Technologies Ltd.) were used to obtain data in Figs. 2(a, d) and Fig. 3. Fluorescence beads (200 nm, F8809, Life Technologies Ltd.) were used to obtain data in Figs. 2(c, f).

Scattering bead phantom: The sample was prepared by embedding fluorescent microspheres in a turbid matrix to mimic light scattering in heterogeneous media. Briefly, 1-μm fluorescent polystyrene beads were mixed with 1-μm titanium dioxide (TiO_2_) particles, which served as scattering centers. The two components were dispersed in deionized water at controlled concentrations and thoroughly vortexed to ensure homogeneous mixing. A small volume of the suspension was then deposited onto a glass coverslip and allowed to air-dry at room temperature, forming a thin scattering layer with fluorescent beads uniformly distributed within the turbid background. The resulting phantom provided a stable fluorescence signal for imaging while introducing controlled optical scattering, enabling quantitative comparison of excitation performance under identical scattering conditions.

#### C. elegans

dRIs247 [Pact-5-mCherry-HA-act-5; Cb.unc-119(+)] was grown to the L4 stage on OP50 and washed off by M9 buffer. Then they were fixed by fixing solution (4% formaldehyde) and rotated at room temperature for 0.5h. The fixed worms were washed with M9 buffer and transferred to a 2% agarose gel pad for imaging. ACT-5, a microvillus-specific actin, was located at the apical site in intestinal cells, which was a biomarker of the intestinal lumen in *C*. elegans. Images of live worm fed with OP50 for 8 days.

Mouse brain sample: C57BL/6J mice were used in the study. Adeno-associated virus (AAV) carrying human synapsin (hSyn) promoter driven tdTomato reporter (pAAV-hSyn-tdTomato) was injected into the mouse brain. After 3 weeks, animals were perfused transcranially with PBS followed by 4% PFA. Harvested brain samples were post-fixed overnight at 4°C in 4% PFA. Brain samples were sectioned into 100-μm thick coronal sections and neurons expressing tdTomato were imaged. All experiments were approved and performed in accordance with Xiamen University Committee on the Use of Live Animals in Teaching and Research guidelines.

## Funding

This work was supported by National Natural Science Foundation of China (62305274, 62475222, 32200919); Natural Science Foundation of Xiamen City, China (3502Z202371001); Fujian Provincial Natural Science Foundation of China (2024J01056); Fundamental Research Funds for the Central Universities (20720240025).

## Disclosures

The authors declare that they have no conflict of interest.

## Data availability

All the data supporting our findings are presented in the main text and the supplemental document. The MATLAB codes are available from the corresponding author upon reasonable request.

## Supplemental document

See Supplementary, Visualization 1, and 2 for supporting content.

## Author contributions

X.W. and Q.S. contributed equally to this work. H.H. conceived the idea. X.W. and Q.S. performed the research. H.H., Q.S., and X.W. analyzed the data. C.K. supported the biological experiments and analysis. X.D. supported the beam design. Q.Z. supported biological analysis. J.D. and H.H. supervised the whole project. H.H. wrote the manuscript with input from all authors. All authors discussed and revised the manuscript.

## Notes

### Competing Interest Statement

The authors have declared no competing interest.

