## Supplementary for "Ultrashort optical-pin excitation for scattering brain imaging"

#### Supporting Information


**Authors**

Xiaobin Weng[1,†], Qiannan Song[1,†], Cihang Kong[2], Xin Dong[3], Qingliang Zhao[4], Jun Dong[1,5,*], and Hongsen He[1,3,5,*]

**Affiliations**

[1] Laboratory of Laser and Applied Photonics (LLAP), Department of Electronic Engineering, Xiamen University, Xiamen 361102, China

[2] Institute for Translational Brain Research, MOE Frontiers Center for Brain Science, Fudan University, Shanghai 200032, China

[3] Department of Electrical and Electronic Engineering, The University of Hong Kong, Hong Kong 999077, China

[4] The State Key Laboratory of Vaccines for Infectious Diseases, Xiang An Biomedicine Laboratory, Center for Molecular Imaging and Translational Medicine, School of Public Health, Xiamen University, Xiamen 361102, China

[5] Fujian Key Laboratory of Ultrafast Laser Technology and Applications, Xiamen University, Xiamen 361102, China

†These authors contributed equally to this work


#### Contents


**Fig. S1:** Experimental setup for optical-pin excitation

**Fig. S2:** Illustration of the controlled parameter for producing the optical pin

**Fig. S3:** Lateral resolution variation of optical pin and Gaussian beam

**Fig. S4:** Phase masks in experiments

**Fig. S5:** Photos of the transparent and scattering phantoms

**Fig. S6:** Focusing property of Gaussian beam and optical pin

**Fig. S7:** Process to create the optical pin

**Fig. S8:** Plane-by-plane comparison of scattering brain images

### 1. Experimental setup for optical-pin excitation

To facilitate practical implementation and broader adoption of optical-pin excitation, we provide detailed experimental configurations illustrating both the primary and alternative realization schemes. As shown in Fig. S1(a), the optical pin can be generated within a standard laser-scanning microscopy platform using a phase-only spatial light modulator (SLM). By encoding a conical phase together with an annular condition on the SLM, the excitation beam is simultaneously shaped into a hollow input and focused with an effectively large physical angle, enabling direct generation of optical pin within a single programmable element. This configuration offers high flexibility and precise control over the excitation geometry.

In addition to the SLM-based implementation, Fig. S1(b) illustrates simplified and power-scalable alternatives based on a conventional axicon. In this scheme, optical-pin excitation can be realized either by illuminating the axicon with an annular beam or by focusing a tightly confined Gaussian beam of small diameter into a large-physical-angle axicon. Both approaches generate a broadened transverse angular spectrum while maintaining a finite cone angle, satisfying the conditions required for optical-pin formation. Compared with SLM-based modulation, the axicon-based implementations are optically simpler, more cost-effective, and capable of handling substantially higher optical power, making them particularly suitable for high-energy excitation and long-term imaging.

Importantly, these alternative configurations can be readily integrated into existing laser-scanning microscopy systems with minimal modification, requiring only the insertion of an axicon or a compact beam-shaping module. The ability to realize optical-pin excitation through multiple experimental routes highlights its robustness and versatility, and underscores its compatibility with a wide range of laser-scanning modalities and experimental constraints.

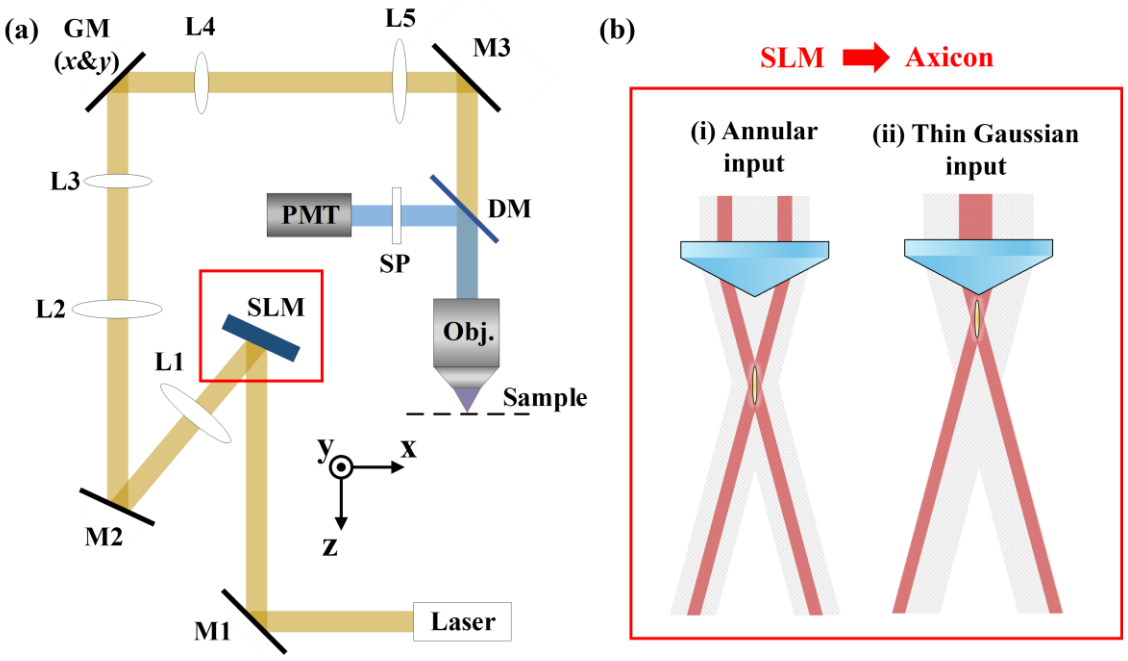


**Fig. S1. Experimental setup for optical-pin excited MPM.** (a) The imaging system is based on a SLM to modulate the light field. M, mirror; L, lens; GM, galvo mirrors; Obj, objective; SP, shortpass filter; PMT, Photomultiplier tube. (b) Alternative approaches by using a single axicon: (i) an annular beam or (ii) a thin Gaussian beam as the input with a high-physical-angle axicon.

#### 2. Illustration of the controlled parameter for producing the optical pin

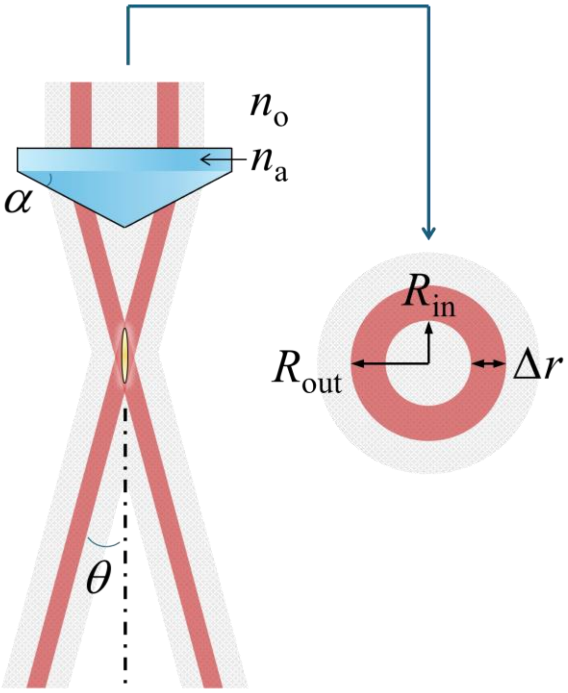


**Fig. S2. Controlled parameter for producing the optical pin.** Schematic definition of the geometric and optical parameters appearing in Eqs. (1–5), including physical/deflection angles, refractive indices, and annular beam dimensions.

#### 3. Lateral resolution variation of optical pin and Gaussian beam

Fig. S3 compares the lateral intensity profiles of optical-pin and Gaussian excitation at different axial positions relative to the focal plane. The figure consists of six panels, with the top row showing optical-pin excitation and the bottom row showing Gaussian excitation. From left to right, the panels correspond to axial positions of -3 μm, 0 μm (nominal focus), and +3 μm. At the focal plane (0 μm), the optical pin exhibits a smaller and more confined central spot than the Gaussian beam. Importantly, when slightly defocused (±3 μm), the optical pin maintains a nearly unchanged lateral spot size and profile, reflecting the persistence of Bessel-type interference over its axial excitation range. In contrast, Gaussian excitation shows pronounced broadening of the lateral spot away from focus, indicating a rapid degradation of lateral confinement under defocus. As a result, the optical pin provides a more stable lateral excitation profile across the axial range, leading to more consistent imaging contrast under practical scanning conditions. Notably, at all three axial positions, the lateral spot produced by the optical pin remains smaller and more confined than that of Gaussian excitation, highlighting its advantage in maintaining uniform excitation conditions throughout the axial section.

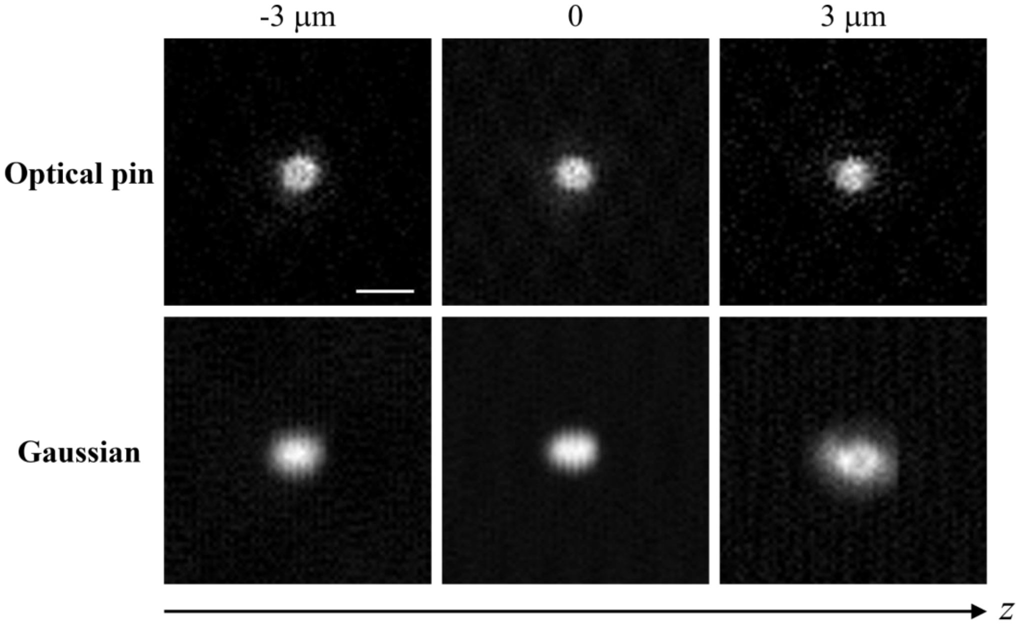


**Fig. S3. Comparison of the lateral resolutions at different axial positions.** Scale bar: 2 μm**.**

#### 4. Phase masks in experiments

Fig. S4 summarizes the phase masks implemented on the SLM to generate different excitation fields in the experiments. The four panels illustrate the progressive shortening of Bessel-type excitation and its contrast with conventional Gaussian focusing. The top-left panel corresponds to a conventional long Bessel beam (B48), generated with a relatively small axicon physical angle, resulting in an extended nondiffracting length of approximately 48 μm. Increasing the physical angle produces a shorter Bessel excitation, shown in the top-right panel (B29), with a reduced axial extent of ~29 μm. The bottom-left panel shows the phase mask used to generate the optical pin (B5). In this configuration, an annular region is selectively extracted from the B29 axicon phase, such that only a narrow radial band contributes to the conical interference. The central disk and outer regions are modulated with an identical linear grating to diffract unwanted light away from the optical axis, effectively implementing annular illumination while preserving the original cone angle. As a result, B29 and B5 share the same axicon angle, but differ in axial extent due to selective angular-bandwidth control. The bottom-right panel shows the Gaussian excitation, implemented using a quadratic (Fresnel lens) phase profile. Together, these phase masks illustrate how progressive control of axicon angle and annular selection enables systematic shortening of Bessel excitation into the ultrashort optical-pin regime, while maintaining a clear distinction from conventional Gaussian focusing. This phase-engineering approach provides a flexible and programmable route for tailoring excitation fields within standard laser-scanning microscopy systems.

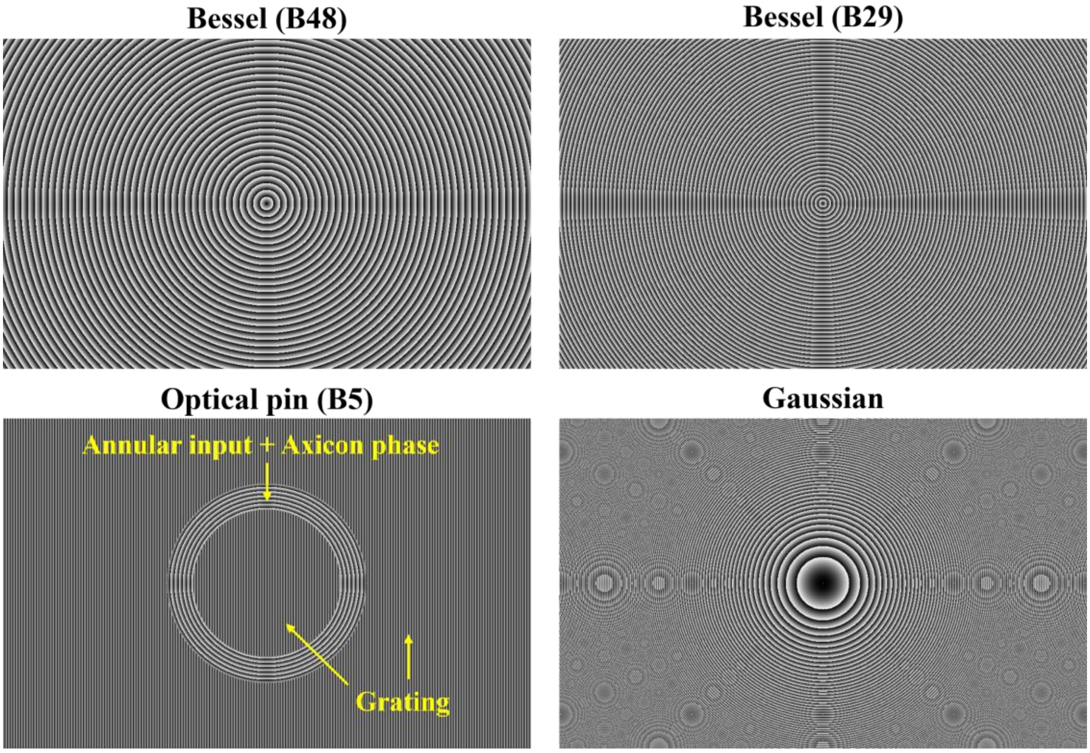


**Fig. S4. SLM phase patterns for nondiffracting Bessel beams, optical pin, and Gaussian beam.**

#### 5. Photos of the transparent and scattering phantoms

Fig. S5 shows photographs of the transparent and scattering bead phantoms used for comparative imaging experiments. The transparent bead phantom was prepared by dispersing fluorescent beads in water without additional scattering particles, resulting in a visually clear sample confined between a glass slide and a coverslip. As shown in Fig. S5(a), the sample appears optically transparent to the naked eye, indicating minimal light scattering and a homogeneous refractive environment. In contrast, the scattering bead phantom contains additional scattering particles, leading to a visibly turbid appearance. As shown in Fig. S5(b), the sample exhibits dark, irregular regions and granular contrast, reflecting the presence of strong optical scattering induced by embedded particles. These two phantoms provide well-defined transparent and scattering conditions, respectively, enabling controlled and direct comparison of excitation performance between Gaussian and optical-pin illumination, as presented in the main text [Fig. 3].

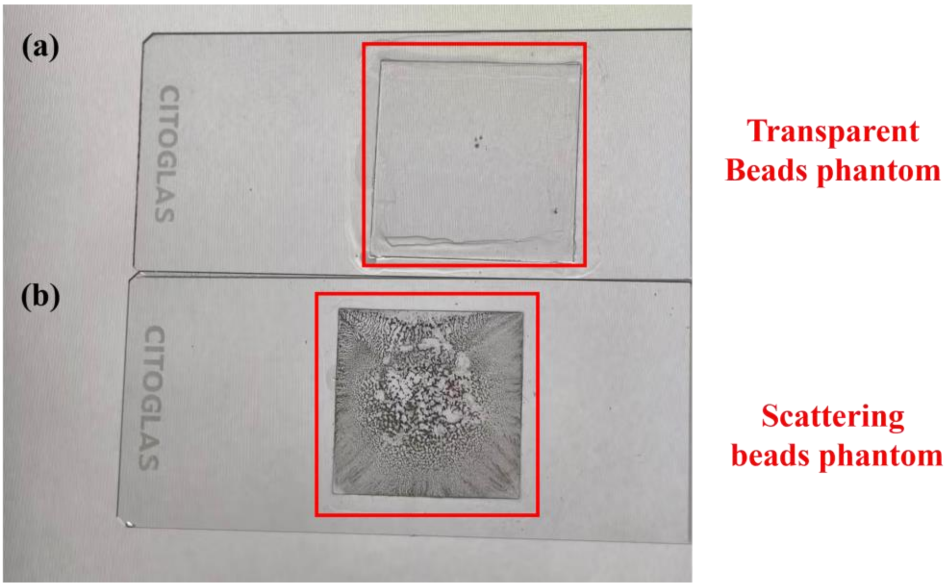


**Fig. S5. Photos of the transparent and scattering phantoms.**

#### 6. Focusing property of Gaussian beam and optical pin

##### Gaussian beam focusing

Conventional laser-scanning microscopy relies on Gaussian focusing, where spherical wavefronts converge toward a single focal point. The focus is formed by a unique phase-matching condition requiring all contributing angular components to arrive in phase. This produces strong confinement under ideal conditions but makes Gaussian focusing intrinsically phase-sensitive, so wavefront distortions or scattering readily disrupt the focus and redistribute energy into the background.

##### Optical pin focusing

Ultrashort Bessel beams are formed by the interference of conical wavefronts associated with a ring-shaped angular spectrum. Rather than converging to a point, these wavefronts intersect the optical axis continuously over a finite range. Axial intensity therefore arises from distributed multi-path interference, making the focusing mechanism less dependent on precise phase alignment at any single position.

##### Response to obstruction and scattering

These distinct interference mechanisms lead to different responses to perturbations. Gaussian focusing degrades rapidly under obstruction or scattering because the focal condition is easily broken. In contrast, Bessel beams mainly lose intensity as contributing paths are reduced, while axial interference persists. As a result, Bessel beams exhibit greater tolerance to obstruction and scattering through statistical path redundancy.

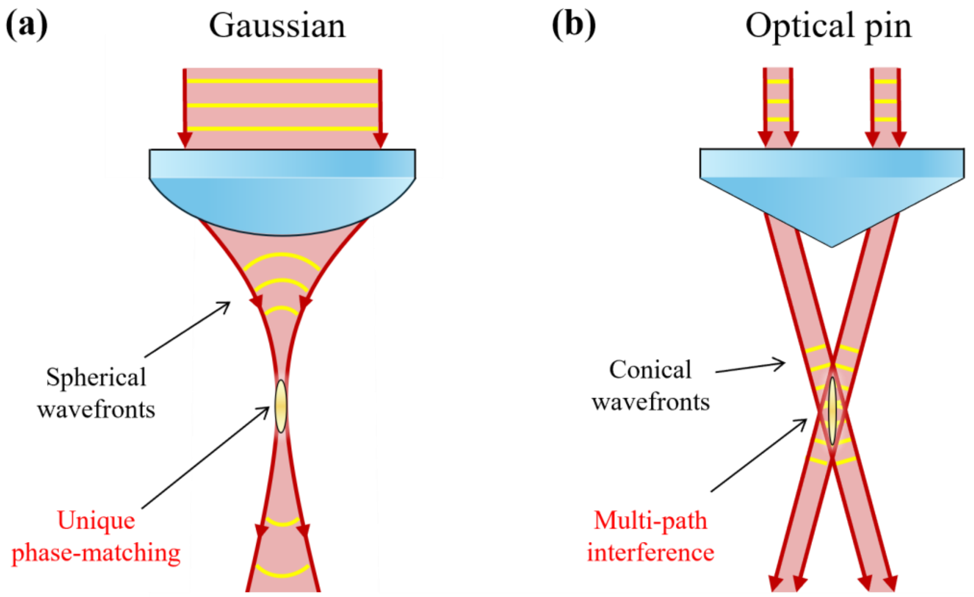


**Fig. S6. Focusing property of Gaussian beam and optical pin.** (a) Gaussian beam. (b) Optical pin.

#### 7. Process to create the optical pin

Fig. S7 is organized into two rows corresponding to the optical-pin (top) and Gaussian (bottom) excitation processes, respectively. In each row, the left column shows the phase masks loaded on the SLM (phase domain), the middle column presents the corresponding Fourier transforms of the phase patterns, representing the transverse angular spectra at the back focal plane of the objective (frequency domain), and the right column displays the resulting axial intensity evolution of the excitation field. For optical-pin generation, the phase mask evolves from a conventional axicon pattern to an annularly selected axicon phase, progressively broadening the transverse angular bandwidth while preserving the conical momentum structure. This spectral evolution is directly reflected in the frequency-domain representation as a transition from a thin annulus to a broadened annular support, which in turn leads to a pronounced compression of the axial interference envelope. In contrast, Gaussian excitation is generated by a quadratic phase profile, producing a centrally concentrated angular spectrum and an axially localized focus whose intensity rapidly degrades away from the focal plane. Together, this video highlights how controlled phase-domain manipulation translates into frequency-domain engineering and ultimately governs the axial structure of the excitation field, providing an intuitive visualization of the optical-pin formation mechanism. **(Visualization 1)**

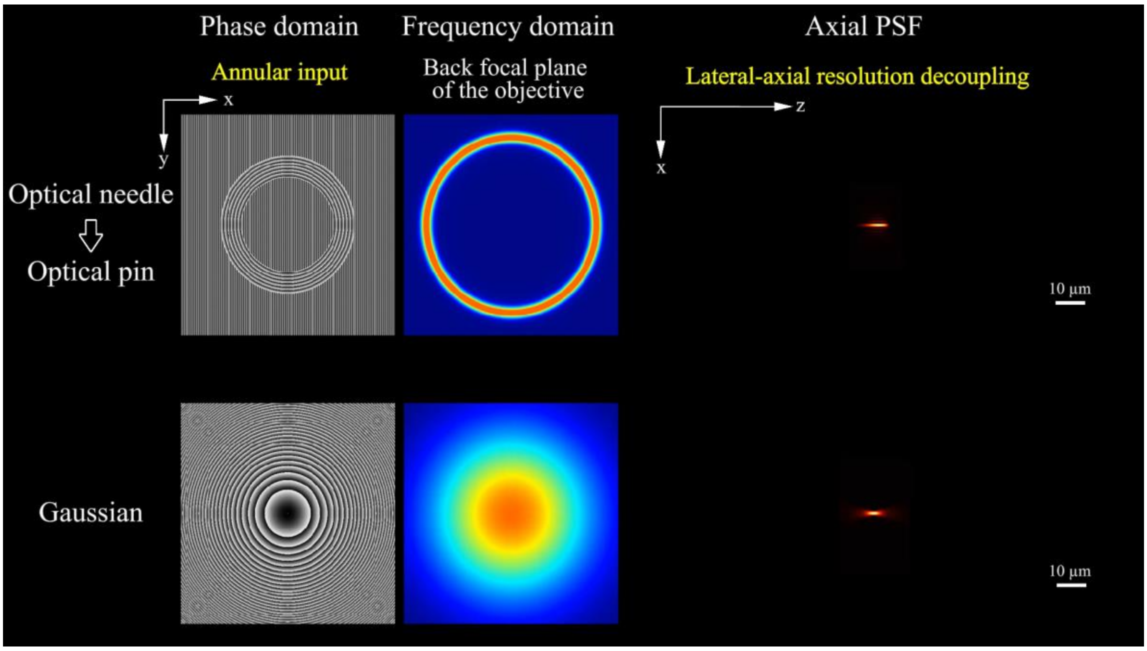


**Fig. S7. Dynamic process of generating the optical pin in comparison with conventional Gaussian beam variation.**

#### 8. Plane-by-plane comparison of scattering brain images

The volume covers an axial range of 20 μm. For each depth plane, the left panel displays the image obtained with Gaussian excitation, while the right panel shows the corresponding image acquired with optical-pin excitation under identical scanning conditions. Across the entire volume, optical-pin excitation consistently provides improved image contrast and clearer visualization of neuronal structures compared with Gaussian excitation. This plane-by-plane comparison demonstrates the overall advantage of optical-pin excitation for volumetric imaging in scattering brain tissue, independent of individual depth planes. **(Visualization 2)**

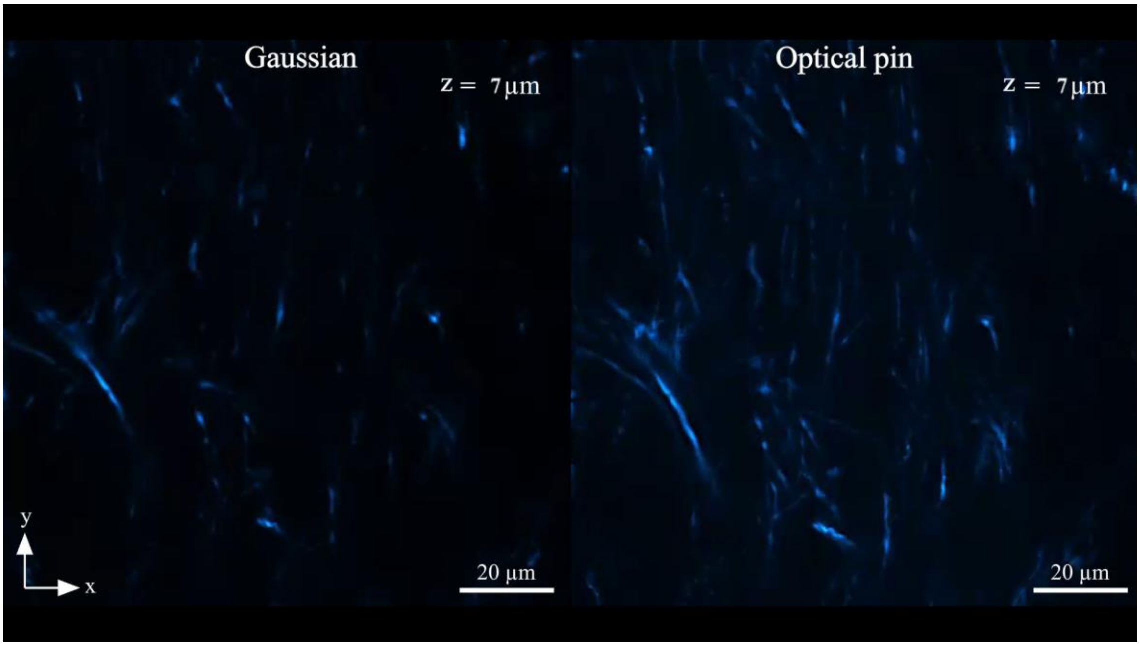


**Fig. S8. Volumetric imaging comparing Gaussian and optical-pin excitation in scattering brain tissue.**
